# Population-level inference for home-range areas

**DOI:** 10.1101/2021.07.05.451204

**Authors:** C. H. Fleming, I. Deznabi, S. Alavi, M. C. Crofoot, B. T. Hirsch, E. P. Medici, M. J. Noonan, R. Kays, W. F. Fagan, D. Sheldon, J. M. Calabrese

## Abstract

- Home-range estimates are a common product of animal tracking data, as each range informs on the area needed by a given individual. Population-level inference on home-range areas—where multiple individual home-ranges are considered to be sampled from a population—is also important to evaluate changes over time, space, or covariates, such as habitat quality or fragmentation, and for comparative analyses of species averages. Population-level home-range parameters have traditionally been estimated by first assuming that the input tracking data were sampled independently when calculating home ranges via conventional kernel density estimation (KDE) or minimal convex polygon (MCP) methods, and then assuming that those individual home ranges were measured exactly when calculating the population-level estimates. This conventional approach does not account for the temporal autocorrelation that is inherent in modern tracking data, nor for the uncertainties of each individual home-range estimate, which are often large and heterogeneous.
- Here, we introduce a statistically and computationally efficient framework for the population-level analysis of home-range areas, based on autocorrelated kernel density estimation (AKDE), that can account for variable temporal autocorrelation and estimation uncertainty.
- We apply our method to empirical examples on lowland tapir (*Tapirus terrestris*), kinkajou (*Potos flavus*), white-nosed coati (*Nasua narica*), white-faced capuchin monkey (*Cebus capucinus*), and spider monkey (*Ateles geoffroyi*), and quantify differences between species, environments, and sexes.
- Our approach allows researchers to more accurately compare different populations with different movement behaviors or sampling schedules, while retaining statistical precision and power when individual home-range uncertainties vary. Finally, we emphasize the estimation of effect sizes when comparing populations, rather than mere significance tests.

## 1 Introduction

Accurately estimating species area requirements is of utmost importance for conservation (Shaffer, 1981; Pe’er et al., 2014), from the individual level to the population level, and especially in light of the increasing human impact on landscapes (Brashares et al., 2001; Dardanelli et al., 2006; Larsen et al., 2008; Nagy-Reis et al., 2021). At the individual level, space-use requirements are typically described by an animal’s home range (Burt, 1943), which is formalized by the probability distribution of the animal’s locations (Worton, 1995). Population-level inference on space-use parameters is also important—both for quantifying the area requirements of a typical organism and for quantifying the effect of covariates, such as species or taxa (Rehm et al., 2018; Habel et al., 2019; Matley et al., 2019; Poessel et al., 2020), sex (Morato et al., 2016; Naveda-Rodríguez et al., 2018; Desbiez et al., 2019; D’haen et al., 2019), body size (Naveda-Rodríguez et al., 2018; Bašić et al., 2019; Desbiez et al., 2019), age (Goldenberg et al., 2018; Averill-Murray et al., 2020; Kays et al., 2020; Mirski et al., 2020), movement characteristics (Swihart et al., 1988; Bowman et al., 2002; Desbiez et al., 2019), conspecific density (Trewhella et al., 1988; Erlinge et al., 1990; Massei et al., 1997), resource density (Massei et al., 1997; Herfindal et al., 2005; Loveridge et al., 2009), habitat or biome (Morato et al., 2016; McBride and Thompson, 2019; Paolini et al., 2019; Tonra et al., 2019), human influences (McBride and Thompson, 2019; Hansen et al., 2020; Rutt et al., 2020; Ullmann et al., 2020), weather (Kay et al., 2017; Matley et al., 2019; Mirski et al., 2020), and season or time (Goldenberg et al., 2018; Roffler and Gregovich, 2018; Bašić et al., 2019; Matley et al., 2019). It has traditionally been the approach that individual home-range area point estimates are input into into conventional analyses (e.g., the sample mean, *t*-test, generalized linear models, etc.) without accounting for their associated uncertainties, likely due to the lack of a suitable alternative (though see Averill-Murray et al., 2020).

Home-range estimation is subject to a number of potential, differential biases that can challenge comparisons across species, behaviors, sampling schedules, tracking devices, or habitats; and only recently have methods been developed to address these issues (Fleming et al., 2015; Fleming and Calabrese, 2017; Fleming et al., 2018, 2019; Noonan et al., 2020; Fleming et al., 2020). Negative biases in home-range estimation can result from less tortuous and less spatially constrained movement behaviors (Swihart et al., 1988; Fleming et al., 2015), finer sampling rates (Swihart and Slade, 1985; Fleming et al., 2015; Noonan et al., 2019*b*), shorter sampling periods (Fleming et al., 2019; Noonan et al., 2019*b*), larger body size (Noonan et al., 2020), and estimating an inappropriate target distribution (Fleming et al., 2015, 2016; Horne et al., 2019). On the other hand, positive biases in home-range estimation can result from over-smoothing of the density function (Worton, 1995; Fleming and Calabrese, 2017) and location error (Moser and Garton, 2007; Thomson et al., 2017; Fleming et al., 2020). These individual-level biases can differ between groups being compared and are expected to propagate into population-level biases, if not dealt with. Furthermore, pragmatic adjustments, such as standardized sampling schedules (Börger et al., 2006) and ‘dilution of precision’ (DOP) values or location-class thresholds (Bjørneraas et al., 2010), will not necessarily avoid differential biases, because home-range estimation biases are a product of both the sampling schedule (or location error) and the movement process. Standardization strategies, therefore, rely on the implicit assumption of the sampled individuals and their tracking devices behaving similarly enough that their biases can be matched by discarding potentially informative data (Winner et al., 2018). This can be acceptable in some cases, but cannot be relied on as a general solution. Instead, statistically efficient estimators that can handle these factors, and can best use all of the data, are necessary to ensure reliable comparisons (Fleming and Calabrese, 2017; Fleming et al., 2020).

To make population-level inferences, researchers and managers traditionally take differentially biased KDE and MCP home-ranged estimates and feed them into general purpose statistical analyses, which assumes that the individual home-range areas are measured exactly (see Signer and Fieberg, 2021, for a thorough description). For instance, a single population would be described by its sample mean, and two populations would be compared with a *t*-test (e.g., Kays and Gittleman, 2001). While the biases of conventional home-range estimators have been studied, and more statistically efficient estimators are now available (Noonan et al., 2019b; Fleming et al., 2020), the impact of ignoring home-range estimation uncertainty on population-level estimates and inferences has not been investigated. More simply, the sample mean of unbiased individual estimates produces an unbiased population-mean estimate, but the sample mean will not achieve minimal variance among all possible population-mean estimators. A more statistically efficient population-mean estimator will down-weight uncertain estimates relative to more certain estimates, in such a way that the estimated mean has a smaller variance, and researchers are not faced with the dilemma of deciding whether or not to include imperfectly sampled individuals when calculating a population average.

In this work we set out to achieve three major goals related to the task of population-level inference on animal home ranges:

1. We demonstrate with empirical data that differentially biased individual home-range estimates propagate into differentially biased population-mean home-range estimates, which is the case for conventional methods that do not account for autocorrelation. In most situations, this bias is negative and sampling dependent, meaning that conventional methods tend to underestimate mean home-range areas (Noonan et al., 2019 *b*).
2. We introduce a novel hierarchical modelling framework for population-level home-range estimation and show that it outperforms existing methods, even in best-case scenarios for the application of conventional methods.
3. We show the problems associated with traditional significance testing on population differences and argue for the estimation of meaningful effect sizes (Sullivan and Feinn, 2012), which we facilitate with a statistically efficient estimator for comparing population home-range areas. Effect sizes provide more information than significant tests and are important for reproducibility (Halsey et al., 2015).

Finally, we conclude with a demonstrative example on multiple frugivore species in the same environment. We implement the introduced methods in the `ctmm R` package `meta` command (version 0.6.0 and later, Fleming and Calabrese, 2015; Calabrese et al., 2016).

## 2 Theory and methods

### 2.1 Effective sample sizes

The effective sample size, *N*, of an individual home-range estimate is the equivalent number of independent and identically distributed (IID) sampled locations required to produce the same quality estimate (Fleming et al., 2019). Biologically speaking, N can also be thought of as the number of observed home-range crossings, as *N* = 𝒪 (*T/τ*), which means that *N* is “on the order of “ *T/τ*, where *T* is the total sampling period and *τ* is the mean-reversion or home-range crossing timescale. When *τ* is large relative to the sampling interval and tracking data are strongly autocorrelated, the effective sample size of a home-range estimate, *N*, can be much smaller than the nominal sample size, *n*, which is the raw number of locations sampled.

As an example, if a tapir crosses its home range twice per day, then, for the purposes of home-range estimation, 12 days of tracking data will be approximately worth as much as 24 independently sampled locations, even if fixes were obtained every second, and over a million locations were recorded. The effective sample sizes of individual home-range estimates can often be small.Even with modern tracking data, Noonan et al. (2019 *b*), in a study of 369 individuals from 27 species, noted that 30% of animal tracking datasets had an effective sample size of less than 30—meaning that many large GPS location datasets were worth less than 30 independently sampled datapoints, for the purpose of home-range estimation. Conventional home-range estimators that assume independently sampled data require hundreds to thousands of observed home-range crossings to produce accurate home-range estimates (Noonan et al., 2019*b*). Moreover, when effective sample sizes are small, home-range estimate uncertainties are large, which are also not accounted for in conventional population-level analyses.

The bias and variance of a home-range estimator is largely a function of the effective sample size, *N* (Noonan et al., 2019*b*). At small-to-moderate effective sample sizes, the most accurate home-range estimators, at present, are based on autocorrelated kernel density estimation, which conditions bandwidth optimization on a fitted autocorrelation model (Fleming et al., 2015). In terms of the autocorrelation estimates, which are the dominant source of bias at small *N*, conventional maximum likelihood and conventional Bayesian methods produce a downward 𝒪 (1/*N*) bias, while residual maximum likelihood (REML) based estimators and the parametric bootstrap can reduce the order of bias to 𝒪(1*/N* ^2^) and 𝒪(1*/N* ^3^), respectively (Fleming et al., 2019). Therefore, to obtain a bias as small as 5%, maximum likelihood and Bayesian methods require on the order of 20 observed range crossings, REML-based methods require on the order of 4−5 observed range crossings, and bootstrapped REML-based methods require on the order of 2−3 observed range crossings. The other important sample size for population estimates is the number of tracked individuals, *m*, and we are also interested in the impact of small *m*.

### 2.2 Hierarchical models

Hierarchical models have long been recognized as providing a natural framework for population-level inference on animal tracking data (Jonsen et al., 2003; Hooten et al., 2016), and here we use them to appropriately weight individual home-range estimates, according to their associated uncertainties, in estimating population-level parameters. Most simply, when modelling the average home-range area of a certain population, it is natural to both consider the individual animal locations to be distributed according to the animal’s home range and to further consider the animals home ranges to be randomly distributed according to their population (Fig. 1). Hierarchical model estimation largely falls into two categories―either fitting the population model to the entire dataset, or first calculating the individual statistics, 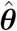, and then fitting the population model to the set of individual statistics. The former is common to Bayesian analyses, while the latter is common to conventional analyses and meta-analyses (Viechtbauer, 2009). Importantly, if 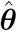 is a *sufficient statistic*―meaning that there is no additional information in the data beyond 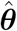 regarding ***θ***―and the exact sampling distribution of 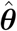 is leveraged, then there is no approximation invoked by the meta-analysis of said statistics (Fisher, 1922). Such is the case with IID Gaussian area estimates, when leveraging their *χ*^2^ sampling distribution. Moreover, even when 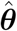 is not a sufficient statistic, maximum-likelihood and Bayesian analyses that fit the population model to the entire collection of tracking data are not exact *per se*, and will generally exhibit both 𝒪(1*/N*) and 𝒪(1*/m*) biases.

**Figure 1:**
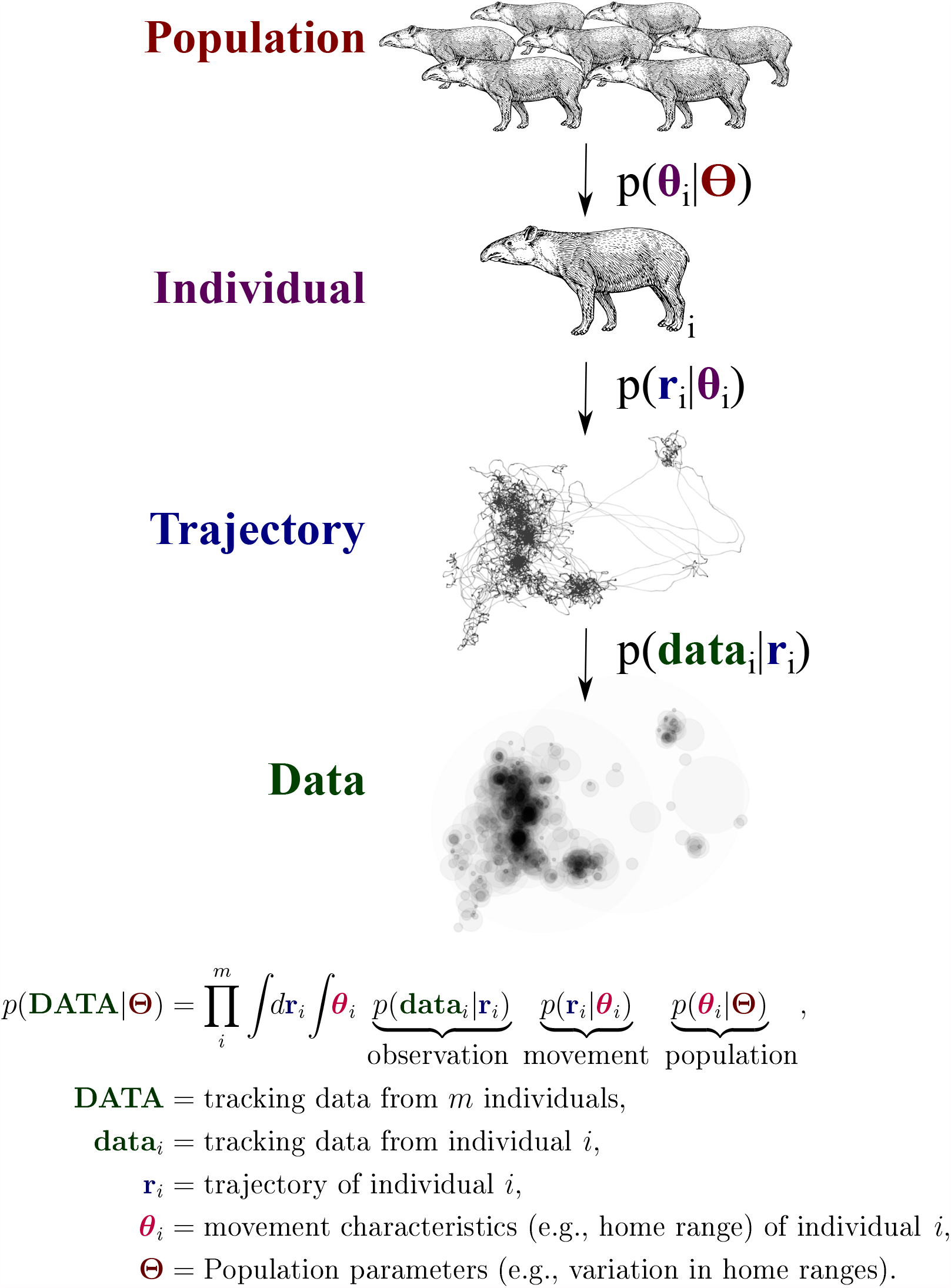
General structure of hierarchical movement models, whereby location data of individual *i* are sparse and erroneous samples of the individual’s unknown trajectory, **r**_*i*_, which is itself the realization of the stochastic process model *p*(**r**_*i*_|***θ***_*i*_), parameterized by movement characteristics ***θ***_*i*_, such as home-range area, mean speed, and diffusion rate. These movement characteristics are, in turn, distributed according to the population model *p*(***θ***_*i*_|**Θ**), which is described by the population parameters **Θ**, such as the population mean of all individual home ranges.

#### 2.2.1 Hierarchical model estimators

In this work we examine the performance of four methods for estimating population-level parameters on animal home ranges―a conventional sample-mean analysis and three proposed alternatives that account for individual uncertainties, including a conventional (normal) meta-analysis, a conventional Bayesian analysis, and a novel *χ*^2^ inverse-Gaussian (*χ*^2^-IG) meta-analysis. All published analyses that we are aware of either neglect home-range uncertainties, and reduce to the sample mean in the absence of covariates, or do not estimate a mean home-range area. So while we refer to the normal meta-analysis and Bayesian analysis as being “conventional”, we are not aware of any pre-existing application or examination of these methods for the task of mean home-range area estimation.

##### Sample-mean analysis

In the conventional sample-mean analysis, we summarize populations by the sample mean of home-range area point estimates, which assumes both large *N* and large *m*. As the sample mean is unbiased, unbiased individual home-range area estimates will be propagated into unbiased population-mean estimates, and vice versa. On the other hand, because the sample mean is unweighted, home-range estimates with higher uncertainty are not down-weighted relative to those with lower uncertainty. As a result, the variance of the population estimate is not minimized when individual uncertainties are heterogeneous. This leaves researchers with the potential dilemma that imperfectly tracked individuals should be omitted from population-level analyses, without clear guidelines on what the threshold of omission should be. In contrast, an optimally weighted mean will produce lower variance population parameter estimates without guesswork, by down-weighting the (small-*N*) uncertain individual estimates. Moreover, because individual home-range area uncertainties are ignored in the sample mean, estimates of population variation will be substantially biased when the number of observed home-range crossings (*N*) is small.

##### Normal-normal meta-analysis

In the conventional meta-analysis, the individual home-range area estimates are modelled as having a normal sampling distribution and the population of home-range areas is also modelled as having a normal distribution. We consider a conventional meta-analysis, because it proposes an easy solution to the challenge of incorporating uncertainties, and is as simple as passing the home-range area point estimates and variance estimates to a single function in `R` (e.g., `metafor`, Viechtbauer, 2009). The normal-normal meta-analytic model is at least somewhat problematic here, as both individual home-range areas and mean home-range areas are positive quantities, which the normal distribution does not respect. A link function could be employed to fix the lower bound, but that approach has two key disadvantages here. First, a link function would not directly produce a *mean*-area estimate, which is our main focus in this work. Second, a link function would give up the unbiased and ‘best linear unbiased estimator’ (BLUE) properties of the normal-normal meta-analysis, whereby unbiased input home-range area estimates will yield unbiased output mean-area estimates, *if* the input uncertainties are correctly specified.

##### Bayesian analysis

In the traditional Bayesian analysis, we consider the marginal likelihood of the entire dataset, given the population model parameters (Fig. 1), with a very weakly informative prior. The conventional Bayesian modelling framework requires us to specify a generative model, but it does not require us to explicitly solve said model, in terms of optimization, integration or density function normalization. Because our likelihood function includes the same conditional density used in the biased maximum likelihood estimation of individuals, we know that these methods will produce large 𝒪(1*/N*) biases, at small *N*, that we want to avoid, for both their posterior predictions of individual ***θ*** and their population-level parameters, **Θ**. To see this, consider the limiting case of a flat (non-informative) population distribution, *p*(***θ*|Θ**), and the opposing case of a singular population distribution. It is then straightforward to show that both cases produce biased predictions and estimates when employing maximum a *posteriori* (MAP) estimation, which is the statistically efficient Bayesian analog to maximum likelihood estimation. Moreover, unless all individuals share the same mean location, increasing the number of individuals, *m*, does not mitigate the small-*N* bias, because increasing *m* only pools a larger number of biased likelihoods.

##### *χ*^*2*^-IG meta-analysis

Finally, we consider a novel meta-analysis framework whereby the individual home-range area estimates are modelled as having a *χ*^2^ sampling distribution, and the population of home-range areas is modelled as having an inverse-Gaussian distribution (Fig. 2). Given the derivations in App. A and included software implementation, this analysis is as simple as feeding our home-range estimates into a single `R` function. However, the distributional assumptions here are far more reasonable than the conventional meta-analysis. In the case of an IID isotropic Gaussian stochastic process, home-range area estimates are sufficient statistics with a *χ*^2^ sampling distribution, and there is no approximation in performing any *χ*^2^-based meta-analysis (versus fitting the population model to the entire dataset). We also find the *χ*^2^ distribution to be a good approximation more generally, for autocorrelated data, which gives the *χ*^2^-based meta-analysis good statistical efficiency when the number of observed home-range crossings (*N*) is small. The choice of an inverse-Gaussian (IG) population distribution will be shown to facilitate good statistical efficiency when the number of sampled individuals (*m*) is small.

**Figure 2:**
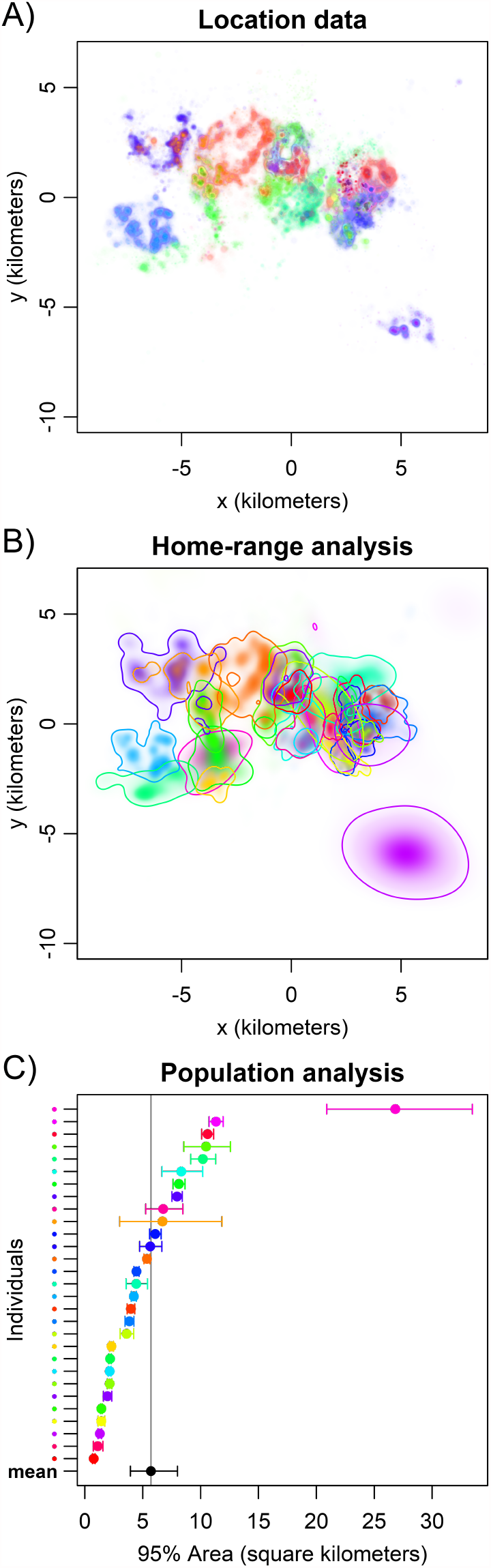
General description of our hierarchical modelling framework, as demonstrated with error-calibrated tracking data on 29 lowland tapir (*Tapirus terrestris*) from the Pantanal region of Brazil. First, animal tracking data (**A**) are used to calculate individual home-range areas (**B**), which are, in turn, fed to our *χ*^2^-IG meta-analysis, here producing a forest plot with the mean home-range area estimate depicted in black (**C**).

#### 2.2.2. Statistical inefficiency of the sample mean

Here we provide some general results on the inefficiency of conventional sample-mean analysis versus any *χ*^2^-based meta-analysis with appropriate limits, in the case of low population variation. Again, *χ*^2^-based meta-analyses achieve increased efficiency over the sample mean by down-weighting uncertain estimates relative to more certain estimates. First, we note that both the sample-mean estimate, 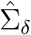, of the mean home-range area, Σ, and the *χ*^2^-based meta-analytic estimate, 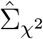, are unbiased if the input home-range area estimates, 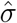, are unbiased. The variances of these estimators are then straightforwardly calculated to be

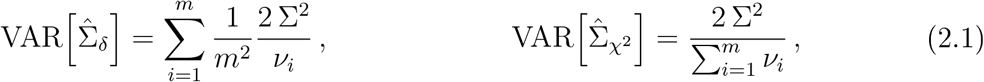

where *ν*_*i*_ represents the degrees of freedom of the *i*^th^ home-range area estimate 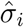, which is given by *ν*_*i*_ = 2*N*_*i*_ for two-dimensional area estimates with effective samples sizes of *N*_*i*_. Next we represent the two variances

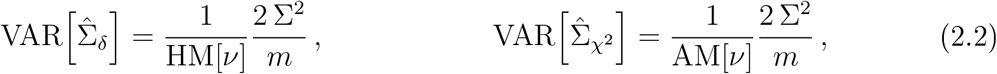

where HM[·] denotes the harmonic mean and AM[·] denotes the arithmetic mean. Pythagorean means are well studied and these two means obey the inequality

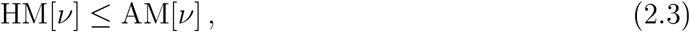

with equality only in the case where all *ν*_*i*_ are equal. Therefore, we have the statistical efficiency inequality

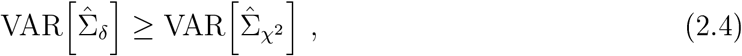

with equality only in the case where all *ν*_*i*_ are equal.

For example, if *ν* = {1, 2, 3, …, 10}, then the *χ*^2^-based 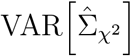 is only 62% of the sample-mean 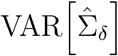, indicating that the sample mean is only 62% efficient for IID Gaussian data, where the *χ*^2^-based estimate is 100% efficient. Moreover, in this case, if the worst two observations are omitted, then the sample mean’s efficiency actually improves from 62% to 81%, whereas the *χ*^2^-based efficiency degrades from 100% to 95%. When using the conventional sample mean, it can be advantageous to discard the worst estimates, whereas in an appropriately weighted analysis, it is advantageous to use all of the data. Finally, we note that there is no importance, in this example, on min(*ν*) = 1. The exact same result follows from *ν* = {10, 20, 30,…,100}. It is the relative differences in individual effective sample sizes that lowers the relative efficiency of the sample mean (Fig. 3).

**Figure 3:**
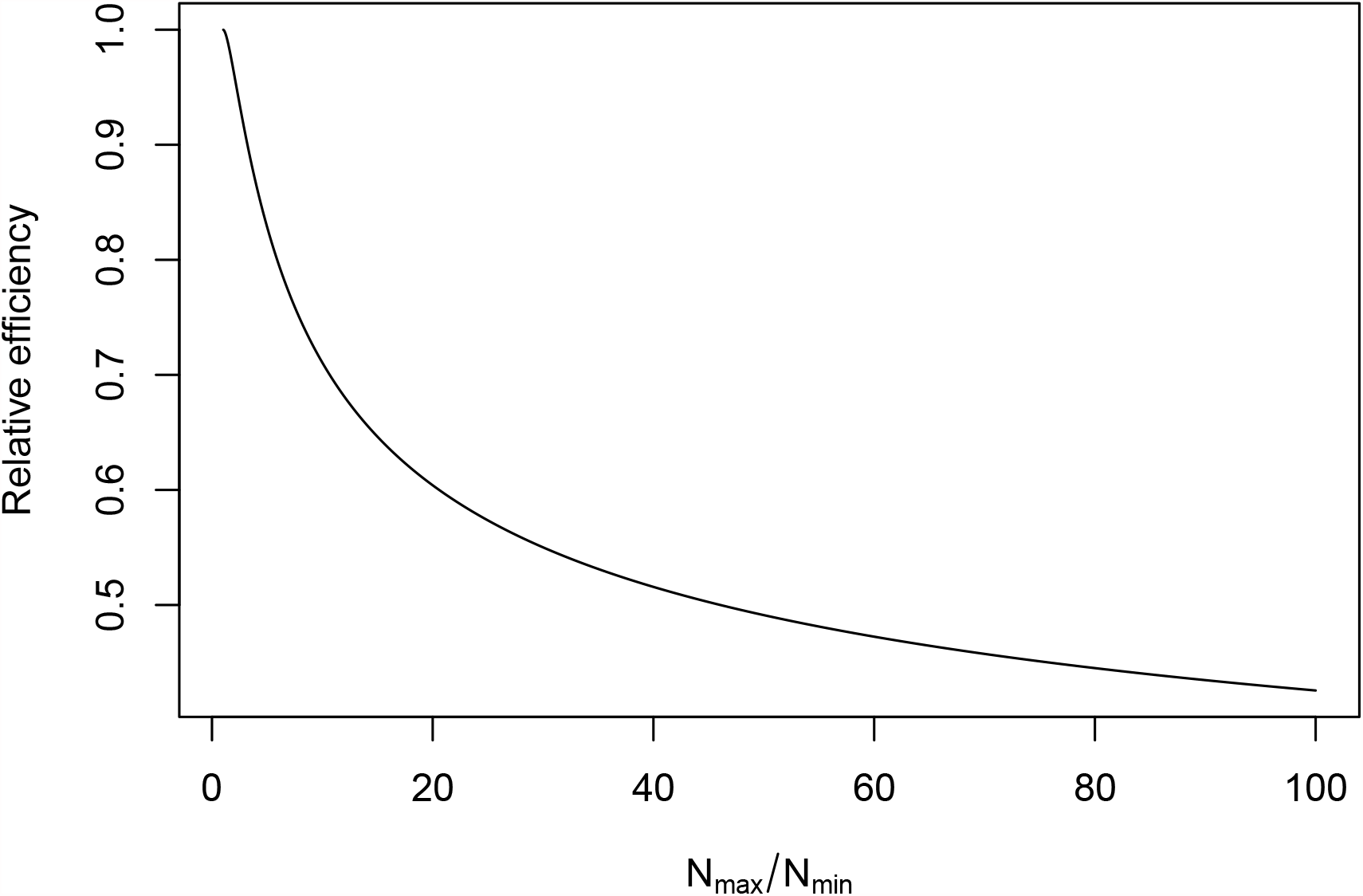
Relative statistical efficiency of the conventional sample mean in comparison to an appropriately weighted mean, when estimating the mean home-range area of a population, and where the effective sample sizes (*N*) of each home-range area estimate are uniformly distributed between *N*_min_ and *N*_max_. All else being equal, the statistical efficiency of the sample mean is worse when there is more variability in the individual effective sample sizes. In general, the unweighted sample mean has a larger variance, which leads to lower statistical power when testing for differences between populations.

#### 2.2.3 The inverse-Gaussian population model

While a *χ*^2^-based meta-analysis can provide good statistical efficiency for small *ν*, the choice of population model and estimators can have a large impact on the statistical efficiency for small *m*, which we now turn our attention to. The mathematically convenient population distribution for a *χ*^2^ sampling distribution is the *inverse-gamma* distribution, as it is a conjugate prior–meaning that the posterior distribution can be calculated in closed form with relative ease. However, the inverse-gamma population distribution is not ensured to produce population mean estimates that fall within the range of the data, and, moreover, produces infinite bias when the shape parameter’s sampling distribution has any support below 1. In contrast, the *inverse-Gaussian* (IG) distribution has a number of important properties that allow us to derive statistically efficient population-level estimates. For instance, in the absence of an hierarchical model, the inverse-Gaussian distribution’s maximum likelihood mean parameter estimate is the sample mean, which is minimum variance unbiased (MVU) (Folks and Chhikara, 1978), and robust to model misspecification.

In App. A we derive a suite of tools for population-level home-range area analysis with a *χ*^2^ sampling distribution and inverse-Gaussian population distribution (*χ*^2^-IG), including debiased estimators for the population mean area, Σ = E[*σ*], inverse population mean area, 1*/*Σ = 1*/*E[*σ*], and square coefficient of variation, CoV[*σ*]^2^ = VAR[*σ*]*/*E[*σ*]^2^, where *σ* denotes a random individual home-range area. We note that having both debiased mean and inverse-mean estimates is important because, as we will discuss, a natural effect size for comparing the home-range areas of populations *I* and *J* is the ratio *R*_*IJ*_ = Σ_*I*_*/*Σ_*J*_, which is the product of Σ_*I*_ and 1*/*Σ_*J*_.

### 2.3 Relevant effect sizes

In the conventional comparative analysis on two populations, *I* and *J*, all individual home-range estimates are calculated each population is summarized by the sample mean of their individual home-range point estimates, 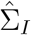 and 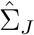; and a *t*-test then determines the statistical significance of any difference in the sample means, 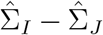 (e.g., Kays and Gittleman, 2001). Here we discourage the overreliance on such *p*-values for several reasons (for further discussion, see Sullivan and Feinn, 2012). Any real pair of different populations will undoubtedly have different mean home-range areas, and the *p*-value is a combined measure of how different two populations are and how much data has been analyzed. As we will show with real data, mean home-range area uncertainties are often relatively large and statistically in-significant *p*-values often do not rule out substantial differences. In their place, we encourage the estimation of relevant effect sizes with confidence intervals. Effect sizes provide more information than *p*-values, are more intuitive, and are important for reproducibility (Halsey et al., 2015).

While the difference between two population-mean home-range areas, Σ_*I*_ *−* Σ_*J*_, is an effect size, it in itself is not informative without the context of what constitutes a large difference. Instead, we consider the ratio

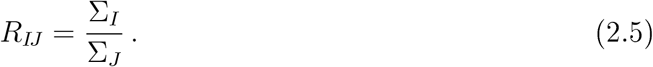

which has a simple biological interpretation that does not require a reference scale to compare to, and also admits unbiased estimators and a relatively simple sampling distribution.

Without loss of generality, let us assume that Σ_*I*_ is greater than Σ_*J*_. If the (two-sided) *α* confidence interval for 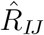 contains 1, then the difference between Σ_*I*_ and Σ_*J*_ is not statistically significant at *p* = *α/*2. However, the difference cannot necessarily be said to be insubstantial unless the confidence interval also does not contain substantial ratios, such as 1.5 or 2. On the other hand, the difference can only be said to be substantial if the confidence intervals do not contain any insubstantial ratios, such as 1.05. What constitutes a substantial effect size is still somewhat subjective, but in reporting effect sizes we avoid conflating data quality with importance.

## 3 Examples

Our examples include three estimator performance comparisons and an empirical analysis demonstration. In our first comparison, we demonstrate with empirical data that the conventional method of taking the sample average of differentially biased individual home-range area estimates results in differentially biased population-mean home-range area estimates. In our second comparison, we contrast the conventional sample mean, conventional normal-normal meta-analysis, and our *χ*^2^-IG meta-analysis on simulated data that are ideal for the sample mean, to demonstrate other advantages of the *χ*^2^-IG framework and serious issues with the conventional meta-analysis (sans link function). In our third comparison, we pit a conventional Bayesian analysis against our *χ*^2^-IG meta-analysis on simulated data, to examine the small-*N* and small-*m* biases of a conventional Bayesian hierarchical estimator. Finally, in our empirical demonstration, we summarize and compare populations (by effect size) in a similar environment, across species and sex.

### 3.1 Estimator performance comparisons

#### 3.1.1. Meta-analytic tapir cross validation

Using our largest dataset of 29 Pantanal lowland tapir with median 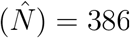, we performed an empirical cross validation analysis to demonstrate the differential bias of conventional population-parameter estimation—whereby biased KDE estimates were fed into the sample mean and compared to more efficient pHREML-AKDE_C_ home-range estimates (Fleming et al., 2019; Fleming and Calabrese, 2017) fed into our *χ*^2^-IG estimator. We chose to cross validate lowland tapir because of the abundance of their data and because they have relatively stable home-range areas (Fleming et al., 2019), which are necessary properties to empirically validate across a wide range of effective sample sizes. To examine the small-*m* bias, we used the entire tracks, where median 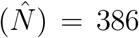, and took random samples of tapir, from 2 to 20 individuals. To examine the small-*N* bias, we used the entire sample of Pantanal tapir (*m* = 29) and incremented the sampling period from 2 to 10 days, with random segments of time sampled from the datasets. Following (Fleming et al., 2019), one day of lowland tapir sampling approximately corresponds to an effective sample size of *N* ≈ 2, and, therefore, 2 to 10 days corresponds approximately to effective sample sizes ranging from 4 to 20.

Given the results of Noonan et al. (2019*b*), we expected the conventional population-mean (Σ) estimates (conditioned on REML-KDE_C_) to increase with increasing *N* and slowly approach the more accurate pHREML-AKDE_C_ estimates for very large *N*. Following Fleming et al. (2019) and Noonan et al. (2019*b*), we expected the pHREML-AKDE_C_ based results to be only slightly underestimated at *N* ∼ 4 (2 days sampling), where they could be improved by bootstrapping (Fleming et al., 2019), which would be too slow for this number of simulations. We expected the reciprocal-mean (1*/*Σ) estimates to exhibit the opposite biases, relative to those of the respective Σ estimates. We expected the conventional population variance estimates to exhibit positive biases that decrease with increasing *N*, as a result of conflation with unaccounted uncertainty.

#### 3.1.2 Meta-analytic simulations

We compared the population parameter estimates of three conditional estimation methods—(1) averaging the point estimates as if they were exact (sample-mean), (2) averaging the estimates with a conventional meta-analytic hierarchical model, which assumes a normal sampling distribution and normal population distribution (normal-normal), and (3) averaging the estimates with our *χ*^2^-IG meta-analytic framework. In our first set of simulations, we incremented the number of observed home-range crossings (*N*) from 1 to 20, with the number of individuals (*m*) set to 100. In our second set of simulations, we incremented the number of individuals from 2 to 20, with the number of observed home-range crossings set to 100. To isolate the biases of the conditional estimators, in all cases we used IID simulated tracking data and Gaussian home-range area estimates, which have ideal statistical efficiency. Furthermore, in each individual meta-analysis, *N* was held constant, so that the conventional sample mean would have its highest efficiency, and other biases could be explored. The population coefficient of variation for the distribution of home ranges was set to one, which is considered to be an intermediate value and was consistent with most of our empirical examples. We performed this analysis both with an inverse-Gaussian population distribution, where our *χ*^2^-IG model is exact, and again with a log-normal population distribution, where our *χ*^2^-IG model is misspecified.

#### 3.1.3 Bayesian simulations

We compared the population parameter estimates of a general-purpose Bayesian estimator to those of our *χ*^2^-IG meta-analytic estimator, when using unbiased REML Gaussian area estimates. We simulated IG distributed home-range areas, and then conditional on those home-range areas we simulated both IID movement processes. The IID tracking data were sampled daily, and would approximate a small canid or deer. The population coefficient of variation for the distribution of home ranges was set to one, which is considered to be an intermediate value and was consistent with most of our empirical examples. We provided our Bayesian estimator with very weak priors that were centered on the truth (App. B), and for output point estimates we considered the mode, median, and mean of the marginal posterior, *p*(Σ). In one set of simulations we incremented the number of observed home-range crossings (*N*) from 2 to 20, with the number of individuals (*m*) set to 100. In a second set of simulations we incremented the number of individuals from 2 to 20, with the number of range crossings set to 100. In this way, we could examine the small sample size biases for both *m* and *N*. For our conditional simulations, we computed 10,000 replicates, while for the much slower Bayesian simulations we computed 1,500 replicates with each having 1,500 draws from the posterior after 1,500 discarded ‘burn-in’ points, after checking for convergence in a number of cases.

### 3.2 Analysis demonstration

#### 3.2.1 Barro Colorado Island frugivore case study

We compared the home-range areas of four species of frugivores—all located on Barro Colorado Island, Panama—including 12 kinkajou (*Potos flavus*), with median 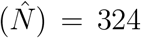, 16 white-nosed coati (*Nasua narica*), with median 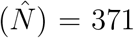, 8 white-faced capuchin monkey (*Cebus capucinus*), with median 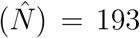, and 8 spider monkey (*Ateles geoffroyi*), with median 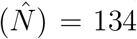. We explored two issues in terms of effect sizes: how home-range areas differed among species and how home-range areas differed between sexes within each species. In all cases we conditioned our *χ*^2^-IG meta-analysis on 95% error-informed pHREML AKDE_C_ home-range area estimates (Calabrese et al., 2016; Fleming et al., 2019; Fleming and Calabrese, 2017; Fleming et al., 2020). Sex differences have been observed for spider monkeys (Campbell,2008), with male home-range areas being larger than those of females, which might also be the case for kinkajous (Kays and Gittleman, 2001). Sex differences have not been observed for coatis (Gompper, 1997), and are not expected for capuchins, because male and female capuchins move together in a social group.

## 4 Results

### 4.1 Estimator performance comparisons

#### 4.1.1 Meta-analytic tapir cross validation

We summarize our lowland tapir cross validation in Fig. 4. We emphasize that these are real data with unknown true parameters, and so we can only assess consistency under resampling. When estimating mean home-range areas (Σ), the conventional KDE estimates increased substantially with increasing sampling period, but did not asymptote enough to match the more accurate AKDE estimates, even when using all of the data. This is more than likely due to many tapir not having the requisite effective sample size necessary for REML-KDE_C_ to exhibit asymptotically efficiency (Noonan et al., 2019*b*). We note that, among home-range estimators that assume independently sampled location data, REML-KDE_C_ is relatively efficient, and other conventional home-range estimators can produce relative biases that are an order of magnitude worse than that of KDE_C_, let alone AKDE_C_ (Noonan et al., 2019*b*). In contrast, when only 2–3 days (*N* ≈ 4–6) were sampled, the pHREML-AKDE_C_ mean home-range estimates had only a slight, negative bias, which could be remedied by bootstrapping (Fleming et al., 2019). However, the bootstrap itself requires repeated simulations, and is too computationally costly to be included in this analysis. As expected, the reciprocal mean area (1*/*Σ) estimates, which are necessary for easily comparing populations, exhibited biases opposite to those of Σ. Finally, both estimators reported extra variation at shorter sampling periods, though this sensitivity was much larger with the conventional estimate.

**Figure 4:**
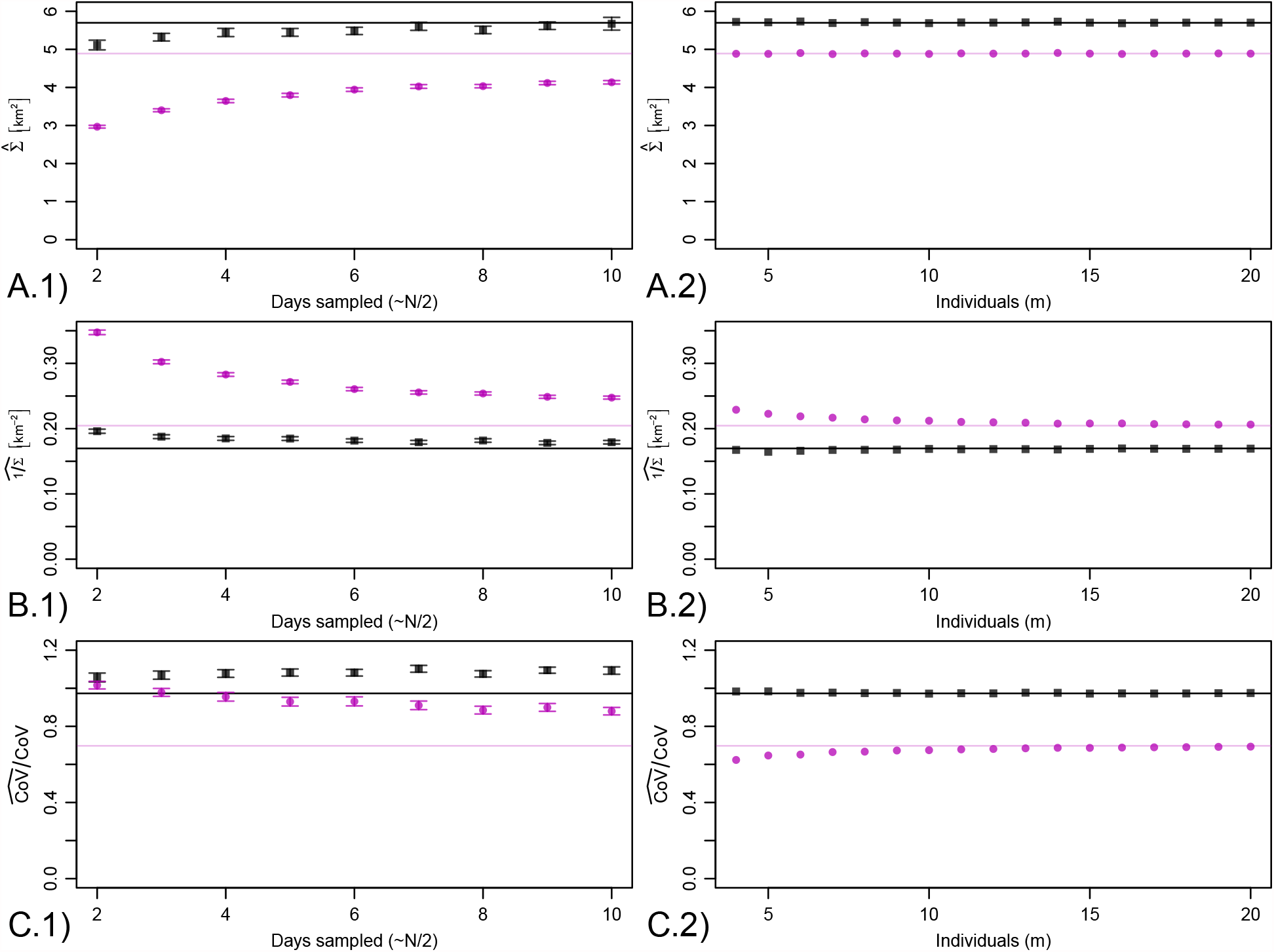
Lowland tapir cross-validation results evaluating two methods for estimating population parameters from tracking data: ignoring individual home-range uncertainties and temporal autocorrelation with sample means of KDE areas (•), versus our *χ*^2^-IG estimators conditional on AKDE home-range areas (▪). On each row of panels a different population parameter is estimated—the mean home-range area (**A**), its reciprocal (**B**), and the coefficient of variation (**C**). In thefirst column, the sampling period is manipulated with all individuals utilized, with the effective sample size (*N*) being approximately twice the number of days sampled. The error bars denote the 95% confidence intervals on the mean point estimates, with 1,000 samples computed. In the second column, the number of individuals (*m*) is manipulated with the full tracks utilized. For reference, the horizontal lines depict the respective point estimates when using all of the data. Overall, the conventional sample-mean KDE estimates were very sensitive to the sampling period and slightly sensitive to the number of individuals sampled.

#### 4.1.2 Meta-analytic simulations

We summarize our meta-analytic simulation comparisons in Fig. 5. Again, these simulations were constructed to be ideal for the conventional sample mean, as in each meta-analysis the number of observed home-range crossings (*N*) was held fixed, which makes the sample mean’s uniform weighting more efficient. Generally speaking, our *χ*^2^-IG conditional estimation framework provided unbiased estimates for all parameters of interest; as expected, the conventional sample mean provided unbiased population mean estimates, but moderately biased estimates of other population parameters; and the conventional normal-normal meta-analysis performed much worse than anticipated, with severe bias at small values of *N*. In our second simulation analysis, with a misspecified log-normal population distribution, the results were almost indistinguishable from the inverse-Gaussian case.

**Figure 5:**
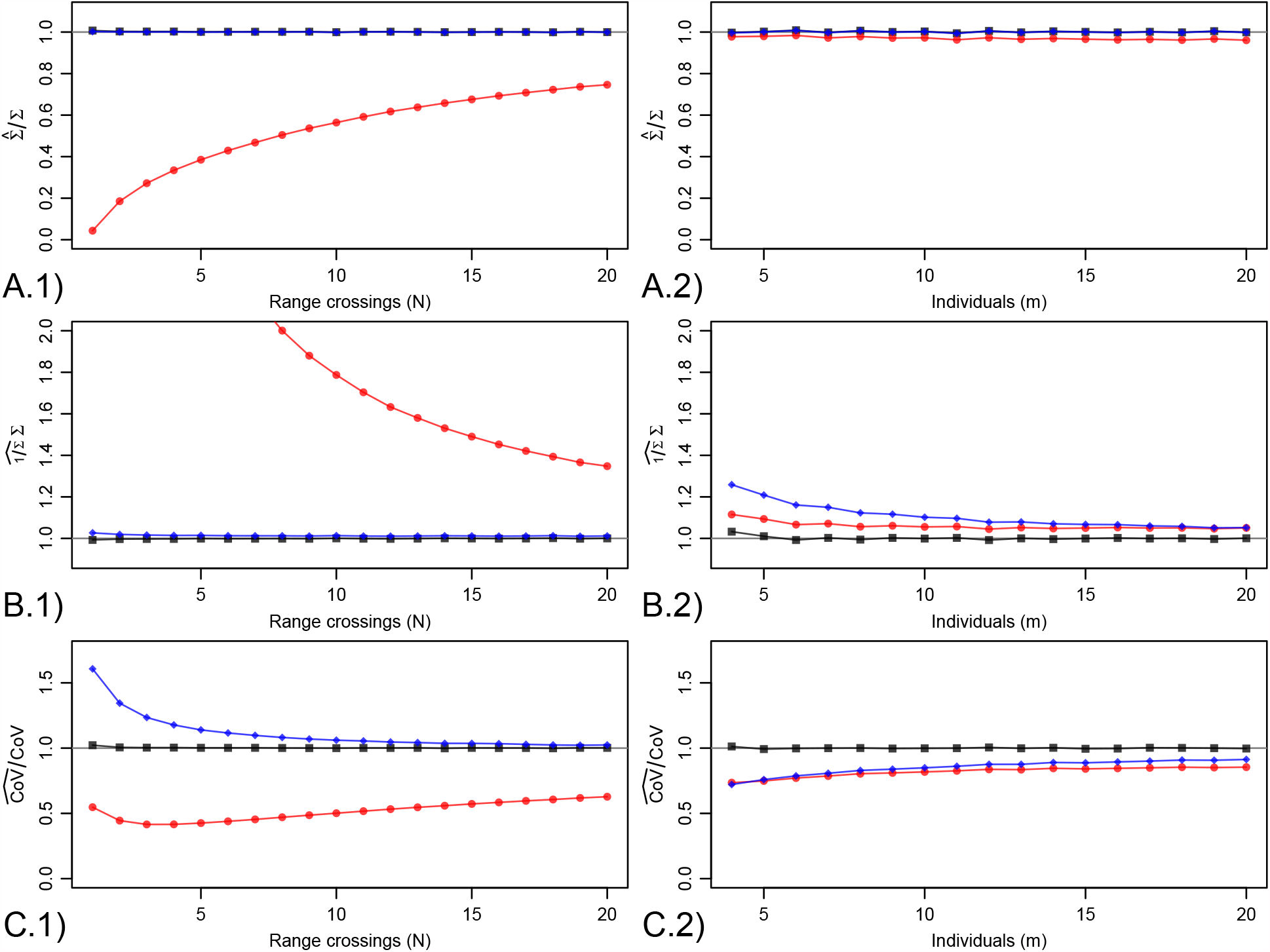
Simulation results evaluating three methods for estimating population parameters from unbiased area estimates: ignoring individual home-range uncertainties with sample means 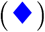, using a conventional normal-normal meta-analytic model to propagate uncertainties 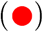, and our *χ*^2^-IG conditional estimator (▪). On each row of panels a different population parameter is estimated—the mean home-range area **(A)**, its reciprocal (**B**), and the coefficient of variation (**C**). In the first column, the number of observed home-range crossings (*N*) is manipulated, while the number of individuals (*m*) is set to 100. In the second column, the number of individuals is manipulated, while the number of observed home-range crossings is set to 100. In general, our *χ*^2^-IG conditional estimators were unbiased, while the other two methods exhibited moderate to severe bias, depending on sample sizes and parameter of interest.

We were initially surprised by poor performance of the conventional normal-normal meta-analytic model, as this method provides approximately ‘best linear unbiased estimates’ (BLUEs) that are exactly BLUE if the variances are correctly specified. However, in retrospect, the variances of a *χ*^2^ process are never exactly known, but are estimated to be proportional to the square of the point estimate. This association causes smaller estimates to be over-weighted and larger estimates to be under-weighted in a normal-normal analysis, which causes the extreme biases depicted in Fig. 5.

Results for *m* = 2, 3 in column 2 of Fig. 5 are not displayed, as they are very much contingent on the chosen model selection criterion and its outcome.If the population variance parameter is supported by model selection, then all *χ*^2^-IG parameter estimates remain relatively unbiased. However, if the population variance parameter is not supported, then the coefficient of variation is taken to be zero and the inverse mean is moderately overestimated. Some degree of model selection is necessary, as the point estimate of the population variance parameter can be in the neighborhood of zero, which would not be selected by any standard model selection criterion, and would cause divergences in both the estimated sampling variances and in the debiased point estimates, if retained.

#### 4.1.3 Bayesian simulations

We summarize our Bayesian simulation comparison in Fig. 6. Generally speaking, our *χ*^2^-IG conditional estimation framework provided unbiased mean area estimates when conditioned on unbiased individual area estimates, as in the previous simulations. In contrast, our Bayesian estimates were far more biased than we anticipated. In terms of observed home-range crossings, we found the small-*N* bias of our Bayesian averages of the anticipated magnitude but in the opposite direction of individual-level maximum likelihood biases. On the other hand, in terms of the number of individuals sampled, we found the small-*m* bias of our Bayesian averages to have an extremely large, positive bias, such that to achieve a reasonable amount of bias, our Bayesian estimator would require more individuals tracked than present in most studies. We tested whether or not this bias was due to a lack of identifiability with the spread of the home-range centers, but this was not the case.Instead, we only found that the small-*m* bias was very similar in scale to the spread of our prior on ∑, even though it was centered on the truth and specified independently of other parameters.

**Figure 6:**
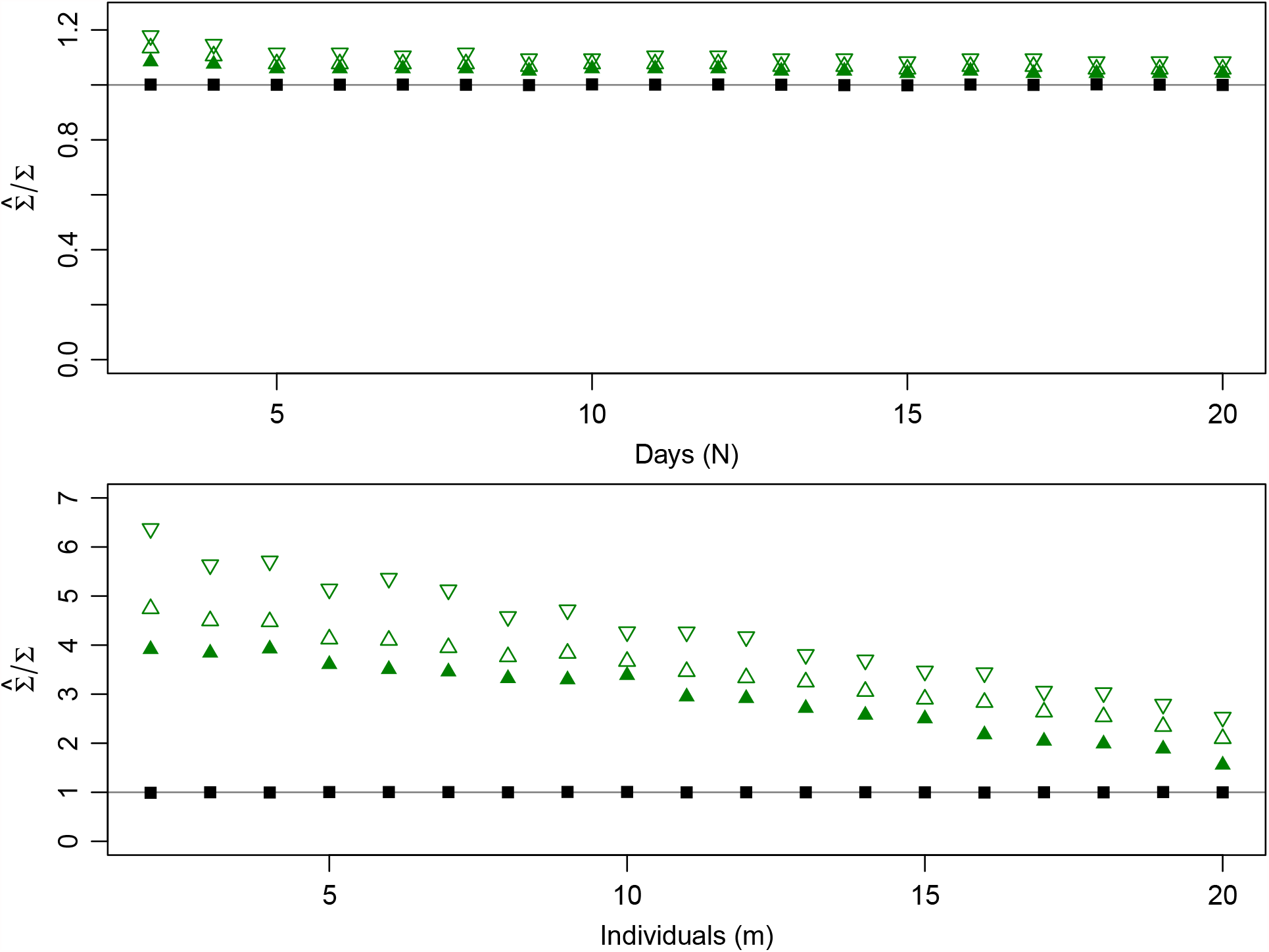
Mean home-range area estimates when using our *χ*^2^-IG conditional estimator on unbiased REML area estimates (▪), and when taking the mode (▽), median (Δ), or mean (▴) of the marginal Bayesian posterior distribution. In the first row, the number of individuals, *m*, is 100, and the observed home-range crossings (*N*) is manipulated. In the second row, the observed home-range crossings is 100, and the number of individuals is manipulated. In general, our *χ*^2^-IG conditional estimators were unbiased when conditioned on unbiased home-range area estimates, while our Bayesian posteriors were much further from the truth than anticipated.

### 4.2 Analysis demonstration

#### 4.2.1 Barro Colorado Island frugivore case study

We summarize the results of our Barro Colorado Island frugivores in Table 1. We found spider monkeys to have the largest home ranges and kinkajou to have the smallest home ranges of the four species, on average, with their mean (95%) home-range areas estimated to be 5.3 (2.6–9.7) km^2^ and 0.3 (0.2–0.4) km^2^, respectively. We could not statistically discriminate the coati and capuchin monkey, and estimated the coati/capuchin ratio of mean home-range areas to be 1.2 (0.8–1.7), which does not rule out a substantial difference. Only in the kinkajou did we find a statistically significant difference between the sexes, where we estimated the male/female ratio of mean home-range areas to be 2.3 (1.5–3.5), which excludes 1, and is both significant and substantial. This test remained statistically significant even if correcting for multiple comparisons. Substantial differences between the sexes could not be ruled out in the other species, due to large uncertainties.

**Table 1:** Results of our Barro Colorado Island frugivore analysis, including the mean home-range area and coefficient of variation (CoV). For reference, a CoV of 1 is considered to be intermediate among distributions on the positive real numbers. ^†^The coefficient of variation for the male Spider monkeys was omale Spider monkeys was not supported by AIC_C_, due to a small sample size (*m* = 3) and relatively large home-range uncertainties, and is, therefore, expected to be underestimated.

|  | mean (km<sup>2</sup>) | CoV |
| --- | --- | --- |
| Spider monkey | 5.3 (2.6–9.7) | 1.0 (0.4–1.6) |
| *Male* | 6.6 (4.7–9.0) | 0.3 (0.1–0.6)<sup>†</sup> |
| *Female* | 4.5 (1.6–10.3) | 1.2 (0.3–2.1) |
| Coati | 1.4 (1.0–1.8) | 0.6 (0.4–0.8) |
| *Male* | 1.3 (0.8–1.9) | 0.6 (0.3–1.0) |
| *Female* | 1.4 (1.0–2.0) | 0.6 (0.3–0.8) |
| Capuchin monkey | 1.1 (0.9–1.5) | 0.3 (0.2–0.5) |
| *Male* | 1.3 (0.8–2.1) | 0.5 (0.1–0.8) |
| *Female* | 1.0 (0.8–1.3) | 0.3 (0.1–0.5) |
| Kinkajou | 0.3 (0.2–0.4) | 0.6 (0.3–0.9) |
| *Male* | 0.4 (0.3–0.6) | 0.4 (0.1–0.6) |
| *Female* | 0.2 (0.1–0.3) | 0.4 (0.2–0.6) |

## 5 Discussion

We have introduced a computationally and statistically efficient hierarchical modelling framework for summarizing and comparing population home-range areas. While we strongly recommend designing studies with larger sample sizes when possible, this framework facilitates population-level inference with as few as 2–3 observed home-range crossings per individual and with a similarly small number of individuals. Importantly, these methods avoid the differential biases inherent in conventional analyses and allow researchers to gain statistical efficiency in using all of their data. In contrast, conventional home-range estimators exhibit downward biases with high sampling rates (Noonan et al., 2020), and even carefully performed data thinning can fail to match these biases across populations (Fleming and Calabrese, 2017). Indeed, data with such high sampling rates require autocorrelation-informed home-range estimators like AKDE. For example, if comparing tapir and jaguar species in the same biome, daily tapir data are more comparable to weekly or monthly jaguar data, for the purpose of home-range estimation (Morato et al., 2016; Fleming et al., 2019), and matching the sampling schedules of these two species can produce wildly different biases from conventional home-range estimators. Here, we have pointed out and demonstrated that these individual-level biases propagate forward into population-level analyses (Winner et al., 2018).

Second, we have proven that, conventional population-level estimators present a second data thinning dilemma to researchers, even when using accurate individual home-range estimates. Conventional population-level estimators perform better when omitting less well-tracked individuals, because certain and uncertain estimates are weighted equally in the sample mean. Indeed, choosing to omit less well-tracked individuals is an extreme form of subjective down-weighting that is not optimized in practice, and would still be outperformed by an appropriately weighted method if it was (Sec. 2.2.2). An appropriately weighted analysis—where uncertain individual home-range estimates are down-weighted relative to more certain estimates—is necessary to produce the best quality population estimates. Our *χ*^2^-IG framework provides said weighting via an hierarchical model.

Third, we recommend and facilitate that researchers comparing populations do so by way of relevant effect sizes, provided with confidence intervals, rather than *p*-values, which are more variable and less reproducible. As we have demonstrated, insignificant differences do not imply insubstantial differences (Sec. 4.2.1). A ratio of mean home-range areas CI of (0.9–2.1) contains 1, which implies an insignificant difference, but it also contains 2, which implies that we are not confident that the difference is insubstantial. On the other hand, a ratio CI of (1.01–1.02) does not contain 1, which implies a significant difference, but it does not contain any substantial difference and, therefore, we are confident that the difference is insubstantial. Effect-size CIs provide a more thorough and meaningful comparison than *p*-values, as with insufficient data, substantial differences can be insignificant, and with abundant data, insubstantial differences can be significant.

### Comparison to other hierarchical methods

While we also considered a conventional (normal) meta-analysis and Bayesian analysis, only our novel *χ*^2^-IG meta-analysis proved to be generally suitable for population-level inference on home-range areas. Conventional meta-analyses also down-weight uncertain estimates, however, as we have shown, their direct application here leads to extreme biases, because of the strong association between home-range area uncertainty estimates home-range area point estimates. We considered the conventional normal-normal meta-analysis without a link function in the hope of obtaining approximate BLUE quality estimates. A link function could improve this method’s performance, but at the cost of the unbiased property and with the additional requirement of having to back-transform the output population-parameter estimates. We recommend that researchers using conventional meta-analytic methods for regression analysis, such as in Averill-Murray et al. (2020), also use an appropriate link function and pay careful attention to their residuals. Otherwise, for the purposes demonstrated here, there is no reason to use any of the conventional analyses over the *χ*^2^-IG estimator.

Finally, our Bayesian analysis produced much larger small-*m* biases that we anticipated, even though we supplied a non-informative prior similar to that suggested by Gelman (2006) for variance parameters, and further assisted our Bayesian analysis by centering each prior on the truth.

### Future analyses

While our *χ*^2^-IG hierarchical model was designed for home-range analysis, it would also provide a natural model for population-level inference on diffusion rates, as they also have an approximately *χ*^2^ conditional sampling distribution. Mean speeds and traveled distances, however, would be better modelled as *χ*-IG (Noonan et al., 2019*a*). Moreover, it would be useful to model both fixed and random effects, especially if the same individuals are being grouped in different populations (e.g., pre- and post-treatment). Fixed-effects might be incorporated via standard IG-regression models (Folks and Davis, 1981), but random effects would require more effort to retain efficiency. For the time-being, regression analyses should be performed with conventional meta-analysis regression methods, and with a carefully selected link function.

## Conclusion

We have shown that accurate population-level home-range estimation requires (1) accurate individual home-range estimates to be fed into (2) an appropriate statistical framework. At present, the most accurate non-parametric home-range estimator is AKDE (Noonan et al., 2019 *b*), which has an associated `R` package (`ctmm`, Fleming and Calabrese, 2015; Calabrese et al., 2016) and graphical user interface (`ctmmweb`, Dong et al., 2017; Calabrese et al., 2021). Upon calculating individual AKDEs, the *χ*^2^-IG meta-analysis that we have introduced here can be evaluated with a single function call, via the `meta` command (Fleming and Calabrese, 2015 Calabrese et al., 2016), which is complete with documentation, `help(meta)`, and example code, `example(meta)`. These combined methods—pHREML autocorrelation estimation (Fleming et al., 2019), AKDE_C_ density function estimation (Fleming and Calabrese, 2017), and *χ*^2^-IG meta-analysis—allow researchers to reap the benefits of using all of their data, avoid differential biases, and achieve greater statistical efficiency than has been possible. Future work will extend these methods to diffusion rates, speeds, and complete movement models, which are a necessary ingredient in the estimation of statistically efficient population density estimates, as well as extended regression analyses.

## Acknowledgments

CHF, WFF, and JMC were supported by NSF IIBR 1915347, MJN was supported by NSERC Discovery grant RGPIN-2021-02758, RK was supported by NSF IIBR 1914928, and ID and DS were supported by NSF IIBR 1914887. This work was partially funded by the Center of Advanced Systems Understanding (CASUS) which is financed by Germany’s Federal Ministry of Education and Research (BMBF) and by the Saxon Ministry for Science, Culture and Tourism (SMWK) with tax funds on the basis of the budget approved by the Saxon State Parliament. This project received funding from the NSF BCS 1440755, the Smithsonian Tropical Research Institute, a Packard Foundation Fellowship (2016-65130) and the Alexander von Humboldt Foundation in the framework of the Alexander von Humboldt Professorship endowed by the Federal Ministry of Education and Research awarded to MCC.

Fieldwork was carried out under IACUC protocol numbers 2014-1001-2017, 2017-0912-2020 and 2017-0605-2020 from STRI, and UC Davis IACUC protocol number 18239.

## A Derivations

### A.1 Hierarchical model for *χ*^2^ distributed estimates

First we approximate our individual home-range area estimates, *s*, as being unbiased sufficient statistics that are (proportionally) *χ*^2^ distributed, conditional on the corresponding true home ranges, *σ*, so that E[*s*|*σ*] = *σ*. This approximation is exact in the case of an isotropic (equal variance in all directions) IID Gaussian process. Our conditional sampling distribution is given by

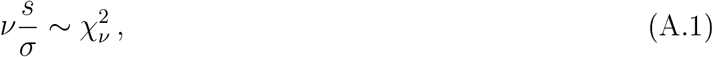

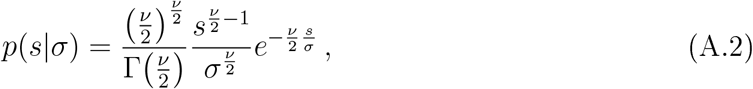

where we treat the degrees of freedom, *ν*, as a known parameter of the individual dataset used to calculate *s*, and VAR[*s*|*σ*] = 2 *σ*^2^*/ν*. For two-dimensional area estimates, *ν* = 2*N*, where *N* is the effective sample size.

For the population-level parameters, we initially consider, most generally, the generalized inverse-Gaussian (GIG) distribution

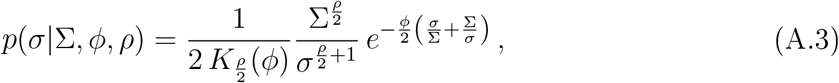

where *K*_*p*_(*ϕ*) is the modified Bessel function of the second kind, and our parameterization here is specifically chosen to make the proceeding integrals concise. The GIG distribution includes, as special cases, the gamma or scaled *χ*^2^ distribution (*ϕ*∑ = 0), inverse-gamma or (scaled) inverse-*χ*^2^ distribution (*ϕ/*∑ = ∞), the inverse-Gaussian (IG) distribution (*ρ* = 1), and the inverse-IG distribution (*ρ* = − 1). We will ultimately settle on the inverse-Gaussian distribution, because it admits statistically efficient parameter estimates, but, for the moment, we continue with the *χ*^2^-GIG calculation.

By straightforward integration, the marginal distribution of the sufficient statistic is given By

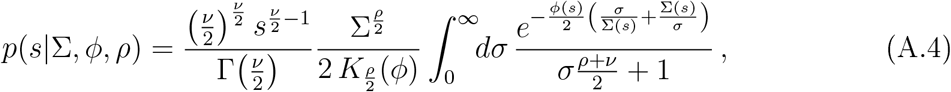

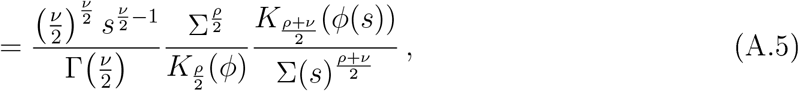

in terms of the functions

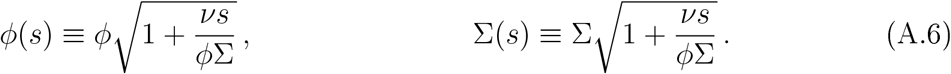

which includes as special cases the *K*-distribution (*ϕ*∑ = 0), the *β*-prime distribution (*ϕ/*∑ = 0), and the scaled *χ*^2^-distribution (*ϕ* = ∞). It is also straightforward to show that 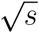 has a half generalized hyperbolic distribution.

There are numerical challenges for evaluating the Bessel functions *K*_*p*_(*ϕ*) at large values of *p* or *ϕ*, which are both necessary in practice. First we consider the log-ratio

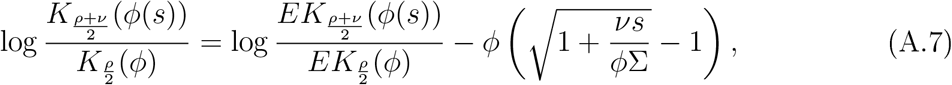

in terms of the exponentially scaled Bessel functions

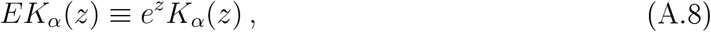

which are implemented in base `R`’ s Bessel functions. We also require, for small arguments, the Padé expansion

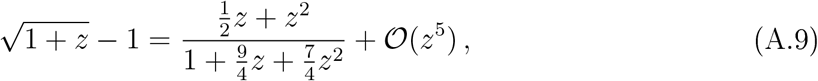

though a Taylor expansion could also suffice, given enough terms. Finally, when *α* is moderate-to-large, *EK*_*α*_ (*ϕ*) can easily exceed the floating point capacity of `xmax` ≈1.797693 × 10^308^ on a 64-bit computer, and base `R` does not implement a log *EK*_*α*_ (*ϕ*) function. This may seem sufficient, but for log-likelihoods we then have a maximum of log(`xmax`) ≈ 709.7827, which is not a particularly large number. Therefore, when base `R`’s *EK*_*α*_ (*ϕ*) approaches `xmax` or returns ∞ at finite values of *α* and non-zero values of *ϕ*, then we resort to Debye’s 1*/α* expansion of log *EK*_*α*_ (*ϕ*) implemented in `R` package `Bessel` (Maechler, 2009), which converges uniformly for both large values of *α* and large values of *ϕ/α*. In our testing, the two implementations agree to within near machine precision when the result is below the `xmax` threshold.

### A.2 Debiased estimators

We now restrict our calculation to the *χ*^2^-IG model, and, following the discussion of Folks and Chhikara (1978), we represent the inverse-Gaussian density function as

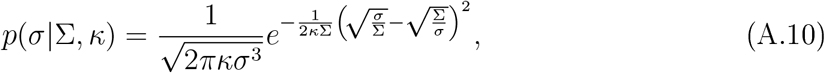

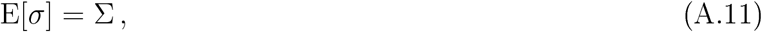

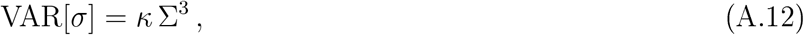

where *κ* = 1*/ϕ*∑ is more convenient for parameter estimation. If the latent variables *σ*_*i*_ are relatively well resolved, or VAR[*s*_*i*_ |*σ*_*i*_]≪VAR[*σ*], we have the maximum-likelihood and minimum variance unbiased (MVU) parameter estimates

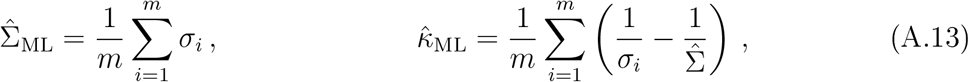

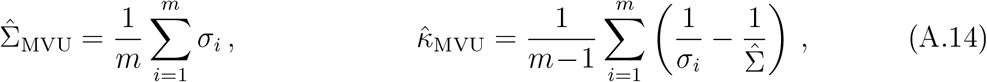

which constitute independent sufficient statistics with sampling distributions

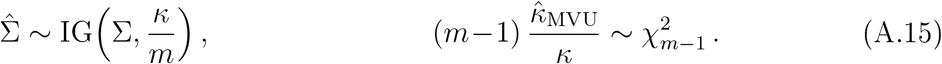

In the opposite parameter regime where the population variation is small compared to individual uncertainties, or VAR[*σ*] ≪VAR[*s*_*i*_ |*σ*_*i*_], then the marginal distribution of *s* reduces to scaled *χ*^2^ and we have the MVU maximum-likelihood estimates

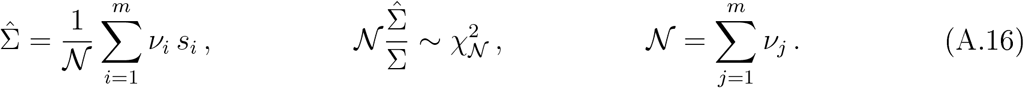

Therefore, by continuity, the maximum-likelihood estimate of the population mean ∑ (argmax of A.5 at *ρ* = 1) must have low bias in general, because the bias vanishes in the opposing parameter regimes of large population variance and small population variance.

On the other hand, the maximum-likelihood estimates of *κ*, VAR[*σ*], STD[*σ*], etc., do have 𝒪(1*/m*) bias. We address this bias approximately, as follows. First, to the maximum-likelihood estimate of *κ*, we include a Bessel correction factor of *m/*(*m* − 1), which is exact for large VAR[*σ*] and makes no difference for small VAR[*σ*]. Second, for population prediction intervals, it is more important to have unbiased estimates of the standard deviation 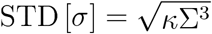. For this task, we note the expectation values

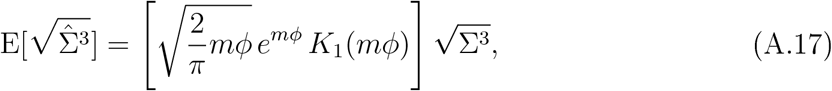

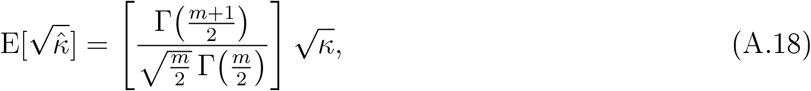

which allow for an approximate first-order bias correction, when dividing by the […] bracketed prefactors evaluated at the parameter estimates.

Finally, we note that the square coefficient of variation is simply given by

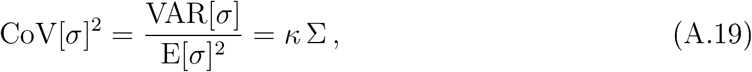

whose plug-in estimator, 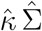, is MVU when VAR[*σ*] is large and the Bessel correction to 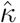 is included.

#### A.2.1 Debiased inverse estimation

Depending on the relative sizes of the estimate uncertainty and population variation, the sampling distribution of 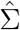 is somewhere between (scaled) chi-squared and inverse-Gaussian. To demarcate this continuum, we introduce the unitless parameter

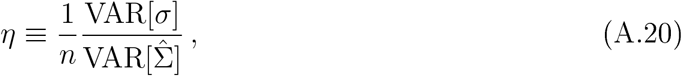

If the population variation is relatively large, then *η* = 1, while if the population variation is relatively small then *η* = 0. To obtain proper limits, we can use *η* to interpolate between the (scaled) chi-square (*η* = 0) and IG(*η* = 1) sampling distributions. It would be more ideal to know the exact sampling distribution of 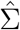, but the maximum-likelihood estimates of likelihood (A.5) are not trivial. With that in mind, the inverse expectation values are given by

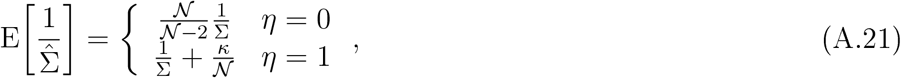

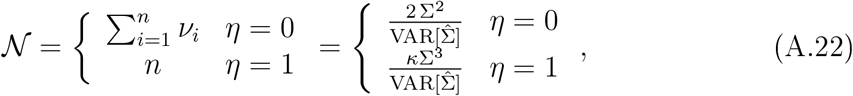

where 𝒩 is a known parameter—the degrees of freedom—in both extremes of *η*, which we can also express in terms of 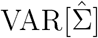. Therefore, after inverting the bias of either relation, by coincidence we have the same inverse estimator

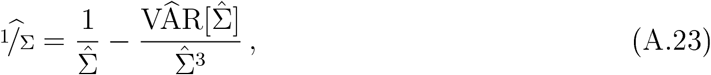

which is MVU at the extremes of *η*, and requires no interpolation for 0 *< η <* 1, where it is at least first-order debiased, as for any unbiased estimator 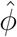

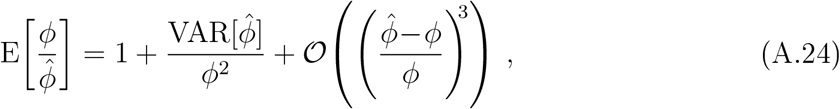

from the Neumann expansion of 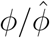 in terms of the error 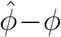.

### A.3 Significance and effect size

Given populations *I* and *J*, we intend to compare the mean home-range areas ∑_*I*_ and ∑_*J*_. Using inverse estimator (A.23), the ratio estimator

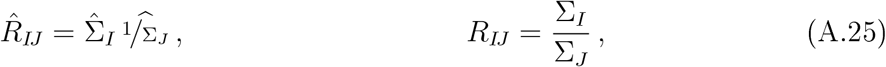

has very little bias. We can closely approximate the sampling distribution of 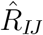 with the scaled *F*-distribution

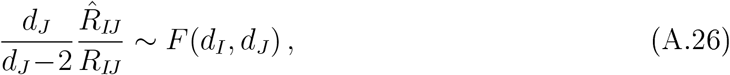

by matching the expectation values and relative variances of the numerator, inverse denominator, and ratio via *d*_*I*_ and *d*_*J*_, in accord with the (scaled) chi-square and inverse-chi-square relations

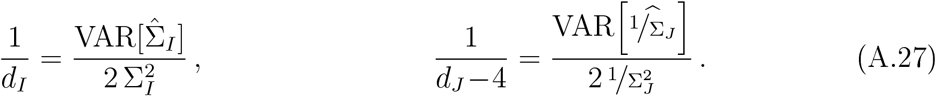

For the numerator, the inverse degrees of freedom has a plug-in estimator that has low bias in both extremes of *η* (A.20):

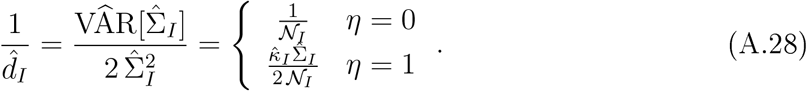

For the inverse-denominator, the relative variance will be slightly underestimated by the delta method, as there are positive corrections to 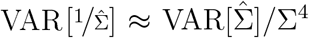 in both regimes of *η*.

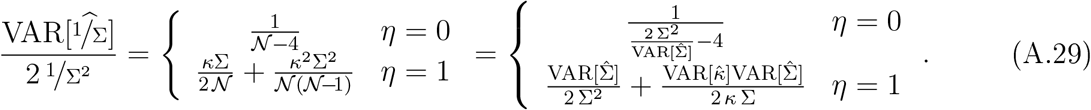

Unfortunately, there is no obvious general expression that bridges the two regimes, so we interpolate the two ends with *η*.

### A.4 Model selection

When home-range estimation uncertainties, 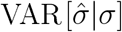, are large compared to the population variation, VAR[*σ*], and especially when *m* is small, then the population variation parameter may not be well supported by the data, and can even be estimated to be zero. Therefore, it is necessary to perform formal model selection between the inverse-Gaussian and Dirac-delta population distributions. Selecting the Dirac-delta population distribution does not imply all individuals have the exact same home-range area, in truth, but that their variation cannot be resolved by the data.

## B Bayesian simulation details

We consider *m* individuals, with the conditional density of the individual *i* with recorded locations **R**_*i*_ being a two-dimensional OU process with individual parameters (latent variables) ***µ***_*i*_, *σ*_*i*_, and *τ*_*i*_ = *σ*_*i*_*/D*_*i*_, where ***µ***_*i*_ is the individual’s mean location, *σ*_*i*_ is the individual’s location variance (in each direction), *τ*_*i*_ is the individual’s home-range crossing or mean-reversion timescale, and *D*_*i*_ is the individual’s instantaneous diffusion rate.

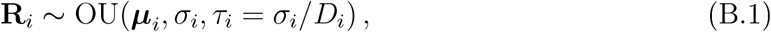

Next we assign the population distributions

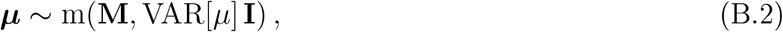

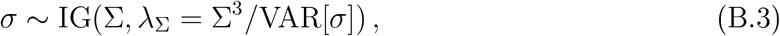

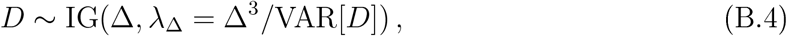

where for simplicity we assign the population variance of ***µ*** to be the same in all directions. The diffusion rates, *D*, are given a distribution independent of the variances, *σ*, because in practice *σ* and *τ* are highly correlated—larger home ranges take longer to cross.

Starting with the population distribution of home-range areas, we assign the population mean home-range area to be a square kilometer and the coefficient of variation to be one.

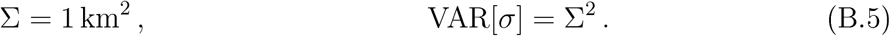

The relative variance in movement (*D*) should be relatively small, as this is a more mechanical or biological parameter, and therefore we set

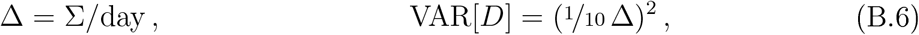

where the mean *D* is such that the average *τ* is approximately a day (given that VAR [*D*] is small). The population distribution of home-range centroids should be a fair amount larger than the individual home-range areas, whereas the population mean location does not matter, and so finally we assign

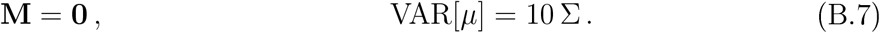

### B.1 Not very informative priors on population-level parameters

For our prior distributions on the population parameters, we use Cauchy and truncated Cauchy distributions that are centered on the truth, but with large scale parameters.

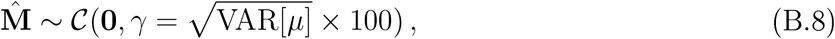

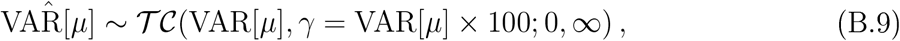

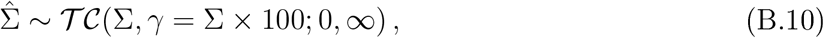

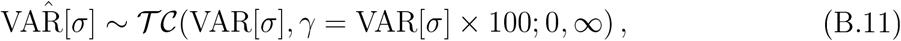

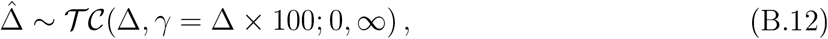

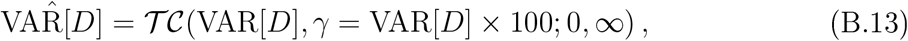

where 𝒞 (*θ, γ*) and 𝒯 𝒞 (*θ, γ*; lower, upper) denote the Cauchy and truncated-Cauchy distributions, with mode *θ* and scale parameter *γ*, and where the truncated-Cauchy distribution also has lower and upper limits specified. The latter choice is similar to the non-informative prior suggested by Gelman (2006) for variance parameters. Therefore, our final prior distributions are given by

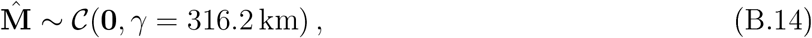

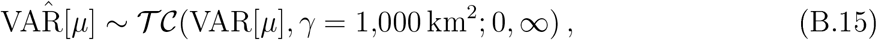

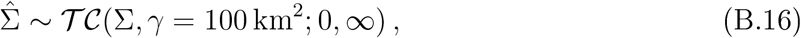

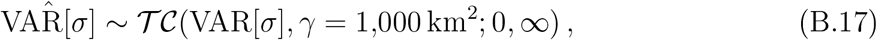

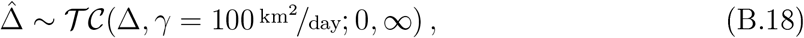

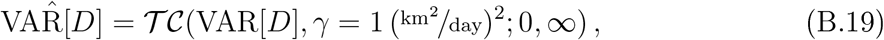

